# Long-term imaging of three-dimensional hyphal development using the ePetri dish

**DOI:** 10.1101/2024.05.18.594613

**Authors:** Oumeng Zhang, Nic Dahlquist, Zachary Leete, Michael Xu, Dean Schneider, Changhuei Yang

**Affiliations:** Division of Engineering and Applied Science, California Institute of Technology, 1200 E California Blvd, Pasadena, CA 91125, USA; Mango Inc, 1314 Westwood Blvd, Los Angeles, CA 90024, USA

## Abstract

Imaging three-dimensional microbial development and behavior over extended periods is crucial for advancing microbiological studies. Here, we introduce an upgraded ePetri dish system specifically designed for extended microbial culturing and 3D imaging, addressing the limitations of existing methods. Our approach includes a sealed growth chamber to enable long-term culturing, and a multi-step reconstruction algorithm that integrates 3D deconvolution, image filtering, ridge, and skeleton detection for detailed visualization of the hyphal network. The system effectively monitored the development of Aspergillus brasiliensis hyphae over a seven-day period, demonstrating the growth medium’s stability within the chamber. The system’s 3D imaging capability was validated in a volume of 5.5 mm × 4 mm × 0.5 mm, revealing a radial growth pattern of fungal hyphae. Additionally, we show that the system can identify potential filter failures that are undetectable with 2D imaging. With these capabilities, the upgraded ePetri dish represents a significant advancement in long-term 3D microbial imaging, promising new insights into microbial development and behavior across various microbiological research areas.

## 1. Introduction

Lens-free on-chip imaging [1] offers a highly compact and optically simple approach to directly capture images of objects. Due to the rapid advancements in CMOS sensor technology that makes them more affordable and higher in quality, this method has become an increasingly viable imaging solution, presenting an economical alternative to conventional imaging methods. The key advantages of lens-free on-chip imaging include an expansive field of view (FOV) that exceeds that of standard microscopes and that is independent of the resolution, a compact and portable form factor, and intrinsic immunity to common optical aberrations due to the absence of lenses. Notable implementations of this technology include the optofluidic [2–6] and digital in-line holographic microscopes [1, 7–11]. However, these methods come with some limitations. For instance, the optofluidic microscope cannot image static samples, which is a significant drawback for observing processes such as cell development. Similarly, the digital in-line holographic microscope strongly depends on the sample sparsity in order for the reconstruction algorithm work properly [12, 13]. To address these limitations, the ePetri dish [14–16] was introduced in 2011. It leverages sweeping illumination angles to achieve sub-camera-pixel resolution without the need to reposition the sample. It not only enables imaging of denser samples but also avoids common noises in coherent imaging, such as the speckles.

However, the ePetri dish has its own limitations; primarily, it produces 2D images - unlike the in-line holography that offers 3D capability. This restricts its ability to render a comprehensive representation of a 3D scene. Further, while the ePetri dish prototype uses a homemade square plastic wall to contain the liquid growth medium for culturing cells, the stability of this medium has not been validated beyond 48 hours, which raises concerns about the long-term integrity of the growth medium for extended biological experiments. Additionally, its adaptability to solid or gel growth mediums is yet to be implemented.

In this paper, we present an upgraded ePetri dish solution that integrates a sealed growth chamber and an advanced reconstruction algorithm, enabling extended microbial culturing and detailed 3D visualization. These improvements significantly extend the capabilities for studying complex microbial growth patterns in 3D, contributing to the field of microbiology and imaging technology.

## 2. Materials and Methods

### 2.1 The ePetri dish and principle of 3D imaging

Figure 1a illustrates the configuration of the ePetri dish designed for extended observation of microbial growth, e.g., bacteria and fungi. This setup uses a randomized 48-LED array to mitigate reconstruction artifacts typically associated with grid LED arrays [14]. One of the primary biological challenges addressed using our setup is the prevention of desiccation of the growth medium over the course of imaging; we designed a sealed plastic culture chamber with a transparent top containing tryptic soy agar. A thin optical film was positioned above the CMOS sensor (Sony IMX183) to create an object-to-sensor gap which enables reuse of the sensor. A representative raw image captured by the sensor is shown in Figure 1b. For this study, *Aspergillus brasiliensis* (*A. brasiliensis*) was chosen as the test organism, initially diluted from a refrigerated spore stock in phosphate-buffered saline, then introduced onto the aforementioned optical film. An optional filter may be inserted between the film and the growth medium to maintain the microorganisms on the film. The complete assembly is commercially available through Mango Inc.

**Fig. 1.**
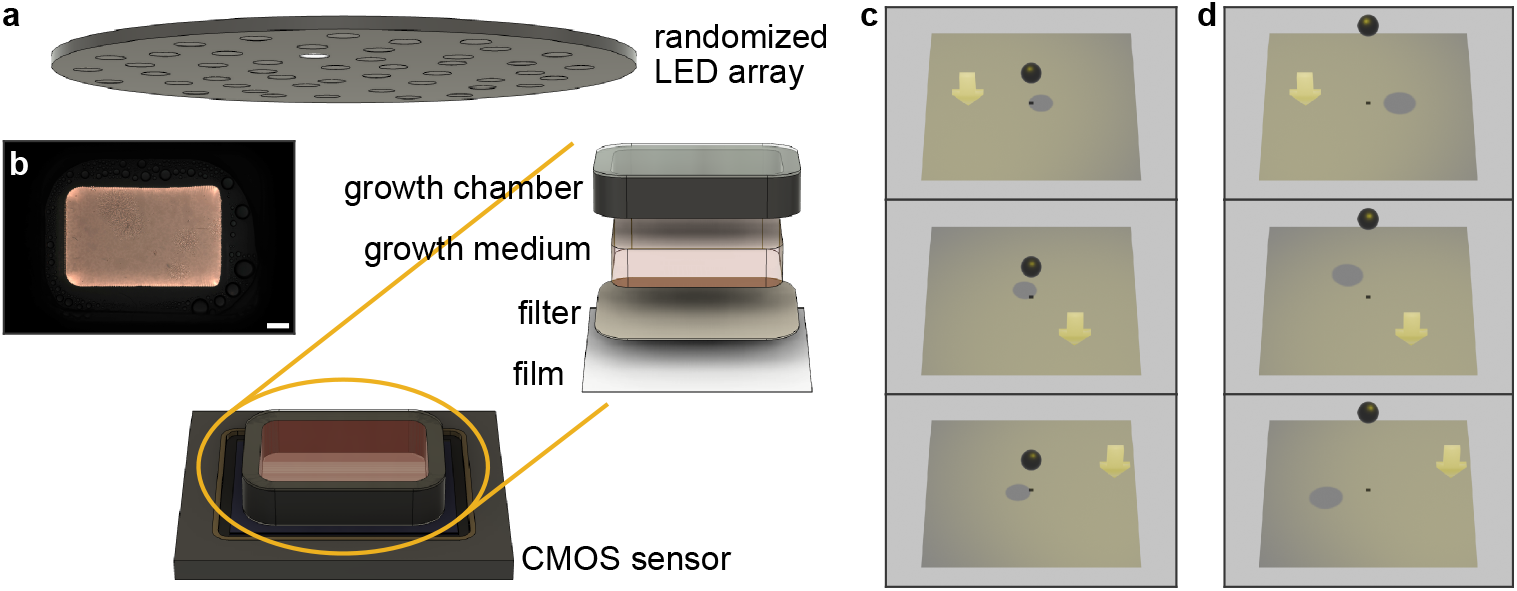
(a) Schematic of the ePetri dish for simultaneous imaging and culture of microorganisms such as bacteria and fungi. The sample is illuminated by a randomized LED array. The culture chamber containing the growth medium is constructed from injection-molded plastic topped with a transparent optical film. An optional filter can be added to inhibit microbial infiltration into the growth medium. A transparent film is placed between the microorganism and the CMOS sensor. (b) Representative image captured directly from the CMOS sensor. Scale bar: 1 mm. (c,d) Principle of three-dimensional imaging using the ePetri dish. The direction of the shadow displacement depends on the LED’s position within the randomized array. (c) An object near the sensor results in a shorter shadow shift relative to the object’s position, compared to (d) an object further from the sensor. The distance between the object and the sensor is determined through the relative positions of the shadows.

The fundamental principle behind the ePetri dish, which enables resolution enhancement beyond the pixel size limitation as demonstrated when it was first proposed [14], leverages the displacement of an object’s shadow when illuminated at tilted angles. By synthesizing multiple shadows cast under various LED orientations, a super-resolved image can be reconstructed. In this work, we further extend the ePetri dish using the proportional relationship between the shadows’ displacement and the object-sensor distance, which allows us to deduce the object’s height by comparing the displacement under different illumination angles (Figures 1c,d). This image formation process is reminiscent of binocular vision in humans, i.e., how human eyes perceive 3D scenes, and also resembles other advanced microscopy techniques such as the Fourier light field microscopy [17], MVR microscopy [18], and the CFAST microscopy [19].

### 2.2 Performance analysis of 3D Imaging using the ePetri dish

We first validate the upgraded ePetri dish’s 3D imaging accuracy and resolution. A Ronchi ruling with 72 lp/mm (equivalent to a line spacing of 6.9 µm) is positioned at a tilt of 46.5°relative to the sensor plane. An algorithm that integrates 3D deconvolution, image filtering, ridge, and skeleton detection (see Section 2.3 for more details) is used to reconstruct the 3D object from the raw images (Figure 2a). The result shows the system’s capacity to resolve individual lines up to a height of 115 µm (Figure 2b), which establishes a sparsity threshold for our samples. For example, for two hyphae positioned 115 µm from the sensor, a minimum separation of ∼7 µm in the *xy* plane is necessary for the ePetri dish to accurately identify both. We also observe a decrease in axial resolution with increased height, with measured reconstruction full width at half maximum (FWHM) values of 5.2 ± 0.7 µm at 15 µm and 14.5 ± 1.2 µm at 75 µm, respectively (Figure 2b, inset). Further, imaging of a tilted clear acrylic plate with three knife-cut lines (Figure 2c,d) validates the accuracy of the reconstruction algorithm; once the sparsity requirements are satisfied, the ePetri dish accurately captures the 3D line structures without noticeable shape distortion up to a height of approximately 2.5 mm (Figure 2e). The measured line tilt matches the ground truth (Figure 2c).

**Fig. 2.**
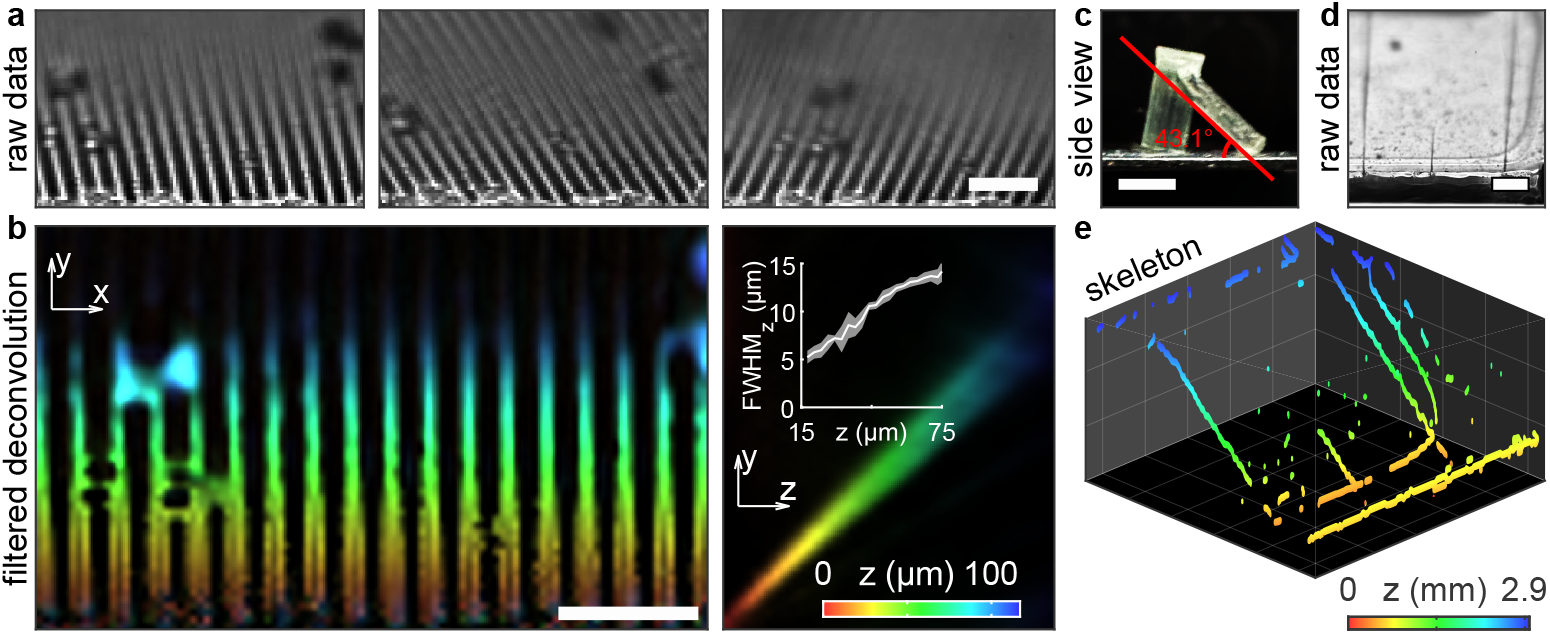
(a) Raw images of a Ronchi ruling positioned at a 46.5°tilt. (b) The *xy* and *yz* views of the 3D reconstruction after image filtering. The inset shows the full width at half maximum (FWHM) along the *z*-direction as a function of height. (c) Side view and (d) raw images of a clear acrylic plate marked with three knife-cut lines, tilted at 43.1°. (e) The 3D reconstructed skeleton of the acrylic plate, with a grid size of 1 mm. Color bar: height; scale bars: 50 µm in (a,b), 5 mm in (c), 1 mm in (d).

### 2.3 Algorithm for reconstructing linear structures from ePetri dish images

We first use a modified Richardson-Lucy deconvolution algorithm [17, 20, 21] to reconstruct the object at varying heights. For sparse samples, the cumulative effect of individual shadow projections at each *z*-plane is assumed to mix incoherently when forming the final image. The corresponding forward imaging model can be approximated as

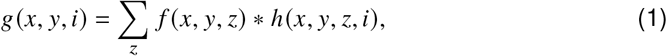

where *g*(*x, y, i*) denotes the image from the *i*th LED (Figure 3a), *f* (*x, y, z*) represents the object, and *h*(*x, y, z, i*) represents the convolution kernel (Figure 3a, bottom insets). To improve computational efficiency, we use *G*(*u, v, i*), *F* (*u, v, z*), and *H*(*u, v, z, i*), the 2D fast Fourier transforms of *g, f*, and *h* to write the iterative deconvolution algorithm as

**Fig. 3.**
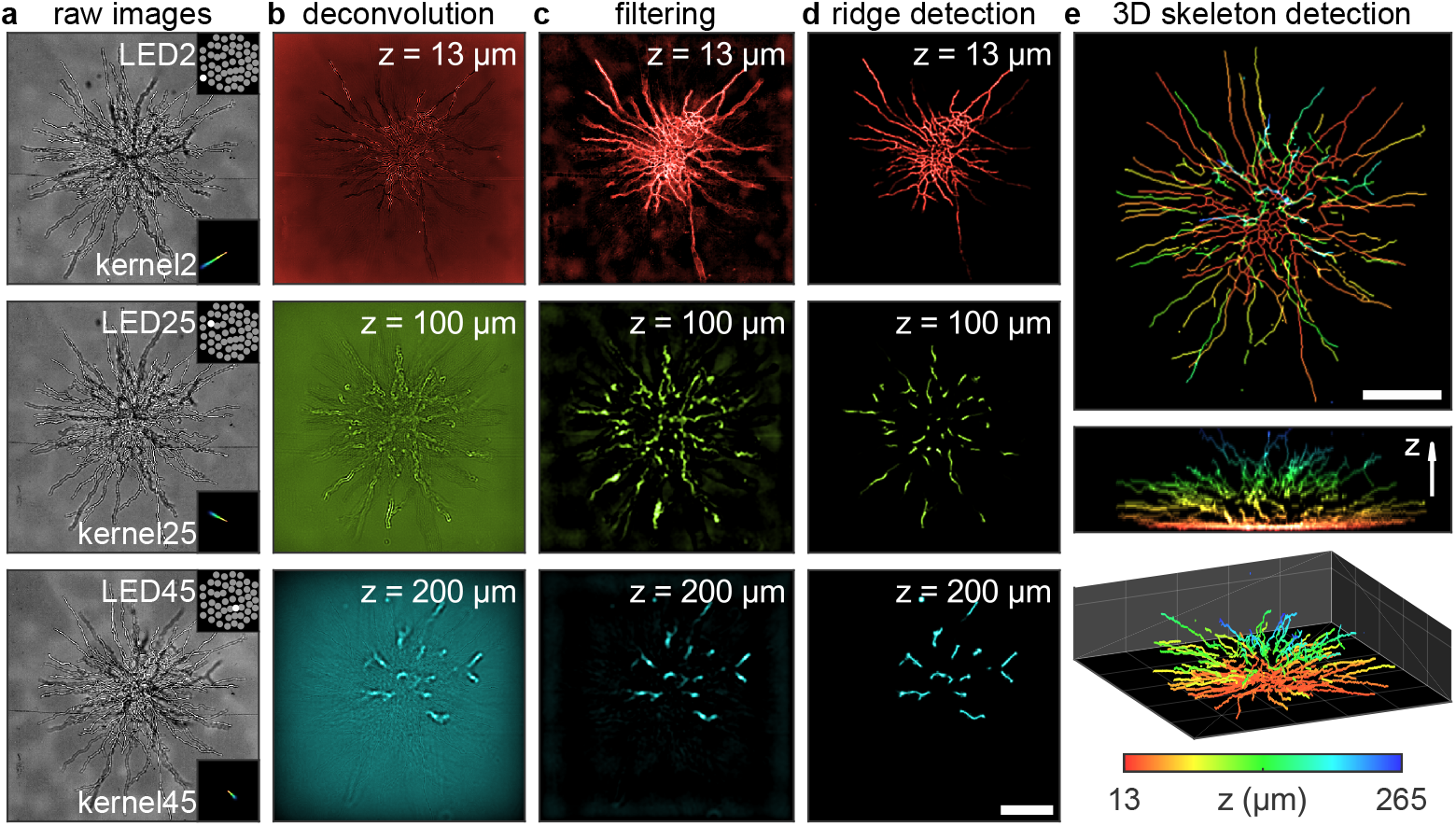
Workflow of the reconstruction algorithm for resolving the hyphae of *A. brasiliensis*. (a) Representative raw images; insets show the active LED and the corresponding deconvolution kernel. (b) Results of deconvolution at *z* = 13 µm, 100 µm, and 200 µm. (c,d) Processed *z*-slice images after (c) image filtering and (d) ridge detection. (e) The *xy, yz*, and 3D view of the 3D skeleton detection results. Scale bar: 200 µm; color bar: height in µm.

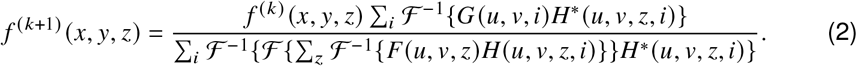

On an Nvidia RTX 4090 GPU, processing a field of view (FOV) of 512×512 pixels with 125 *z* slices (Figure 3b) requires approximately 105 seconds.

We next use a high-pass filter to remove low-frequency background variations. Given the visual characteristics of hyphae close to the sensor (Figure 3b, 13 µm), which exhibit dark cores surrounded by bright edges, we use the MATLAB function “imclose” to perform morphological closing. After filtering, the images of hyphae are significantly clearer (Figure 3c). However, we still observed a decrease in reconstruction quality with increased distance from the sensor due to the approximately linear resolution degradation. To enhance the visualization, we leverage the inherent sparsity of the structure of the hyphae. Similar to how single molecule localization microscopy [22–24] can localize the position of a point source with much higher resolution from a diffraction limited spot, it is possible to identify the hyphae with improved spatial resolution from the blurred images, owing to its linear nature. To achieve this, we implement a Hessian-based mixed 2D (near-sensor *z* slices) and 3D (throughout the volume) ridge detection algorithm [25] to identify the ridges within the reconstructed volume (Figure 3d). To optimize processing speed, closed-form parallel eigen-decomposition is applied [26]. Finally, a 3D skeleton detection algorithm [27,28] is implemented to visualize the hyphae network as shown in Figure 3e. The filtering, ridge detection, and skeleton detection processes require 0.4, 5, and 2.6 seconds respectively, to execute on an AMD Ryzen 9 7900X CPU for the aforementioned FOV.

## 3. Results and Discussions

### 3.1. Long time-scale, large field of view imaging of A. brasiliensis hyphae development

We demonstrate the long-term imaging capabilities of the ePetri dish by culturing *A. brasiliensis* at a controlled temperature of 25°C. This experiment was conducted without a filter. We were able to continuously track *A. brasiliensis* growth over a seven-day period, with no visible degradation of the growth medium. Tracking five initial spores across an FOV of 5.5 mm × 4.0 mm (Figure 4a), the initiation of hyphae growth was recorded at 36, 61, 69, 60, and 73 hours post-inoculation, ordered from the topmost to the bottommost spore. The system achieved a depth of field of approximately 500 µm in this capture.

**Fig. 4.**
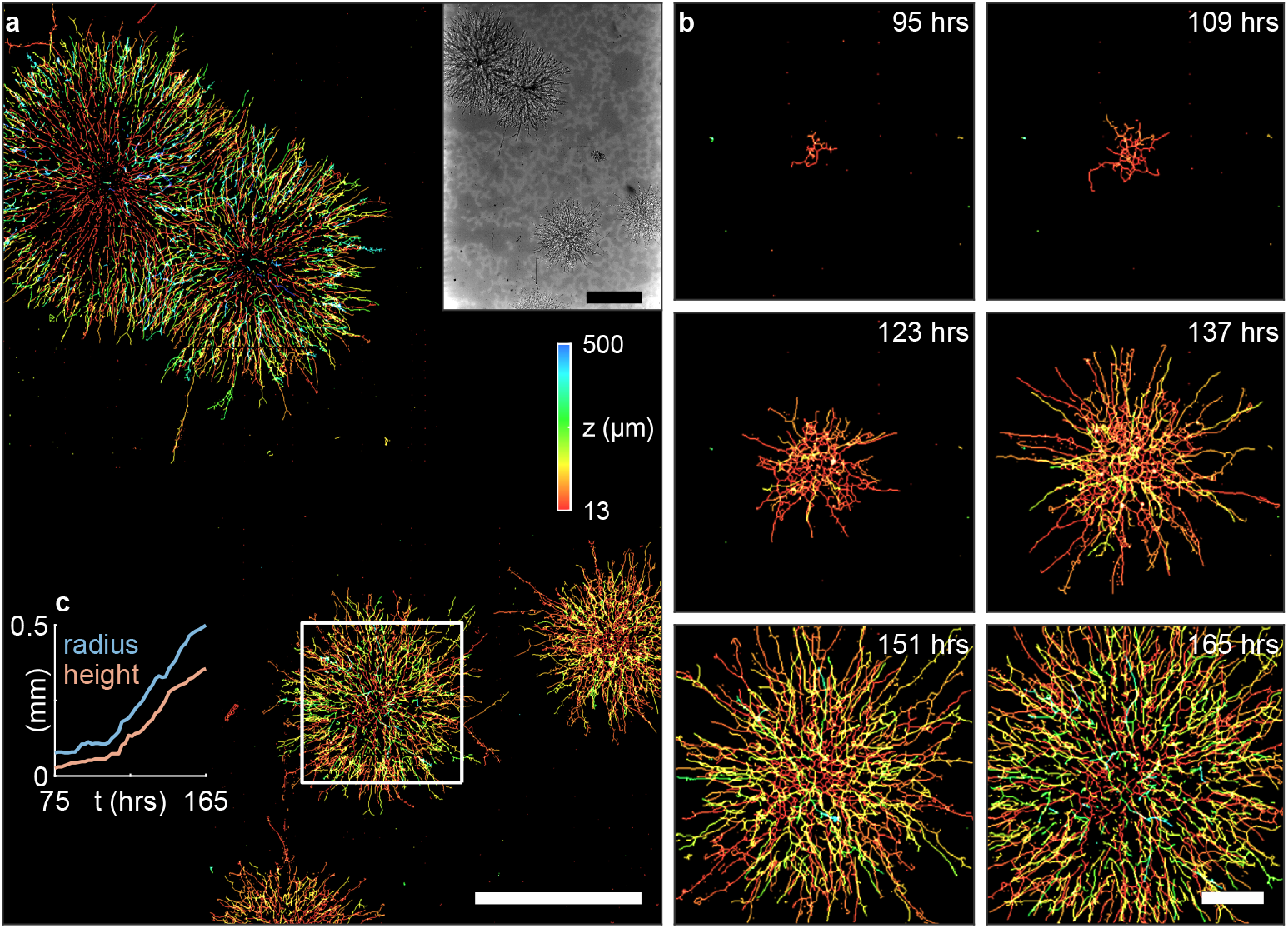
Time-lapse imaging of *A. brasiliensis* development. (a) The entire field of view captured at the end of the time lapse. Inset shows one of the raw images. Scale bar: 1 mm. (b) Boxed area in (a) at multiple time points. Scale bar: 200 µm; color bar: height in µm. (c) Radius and height of the hyphal network in (b) over time.

Figure 4b showcases the development of one particular colony. By tracking both the radius and height of the hyphae network (Figure 4c), we observed a pattern of radial expansion. Interestingly, the vertical growth was slightly slower compared to the radial; at the 165-hour mark, the colony measured a radial expansion of 499 µm and a vertical growth of 355 µm.

### 3.2 Detection of filter failure

A key advantage of the ePetri dish, when compared to coherent on-chip imaging methods like in-line holography, is its robustness to noise from weak-scattering dense objects, such as filters with a slight refractive index mismatch compared to the growth medium. Here, we investigated the growth dynamics of *A. brasiliensis* in the presence of a filter, with the incubation temperature raised to 32.5°C to accelerate growth. The filter proved largely effective in maintaining most of the hyphae close to the CMOS sensor (Figure 5a), with only the tip of a single hypha extending up to 35 µm from the sensor. This segment of the hyphae is undetectable in the super-resolved image when processed using a 2D reconstruction algorithm, as indicated by arrows in Figure 5a. In a separate experiment, we observed early-stage hyphal penetration through the filter into the growth medium (Figure 5b), suggesting a potential compromise in filter integrity which could not be assessed through 2D reconstruction algorithms.

**Fig. 5.**
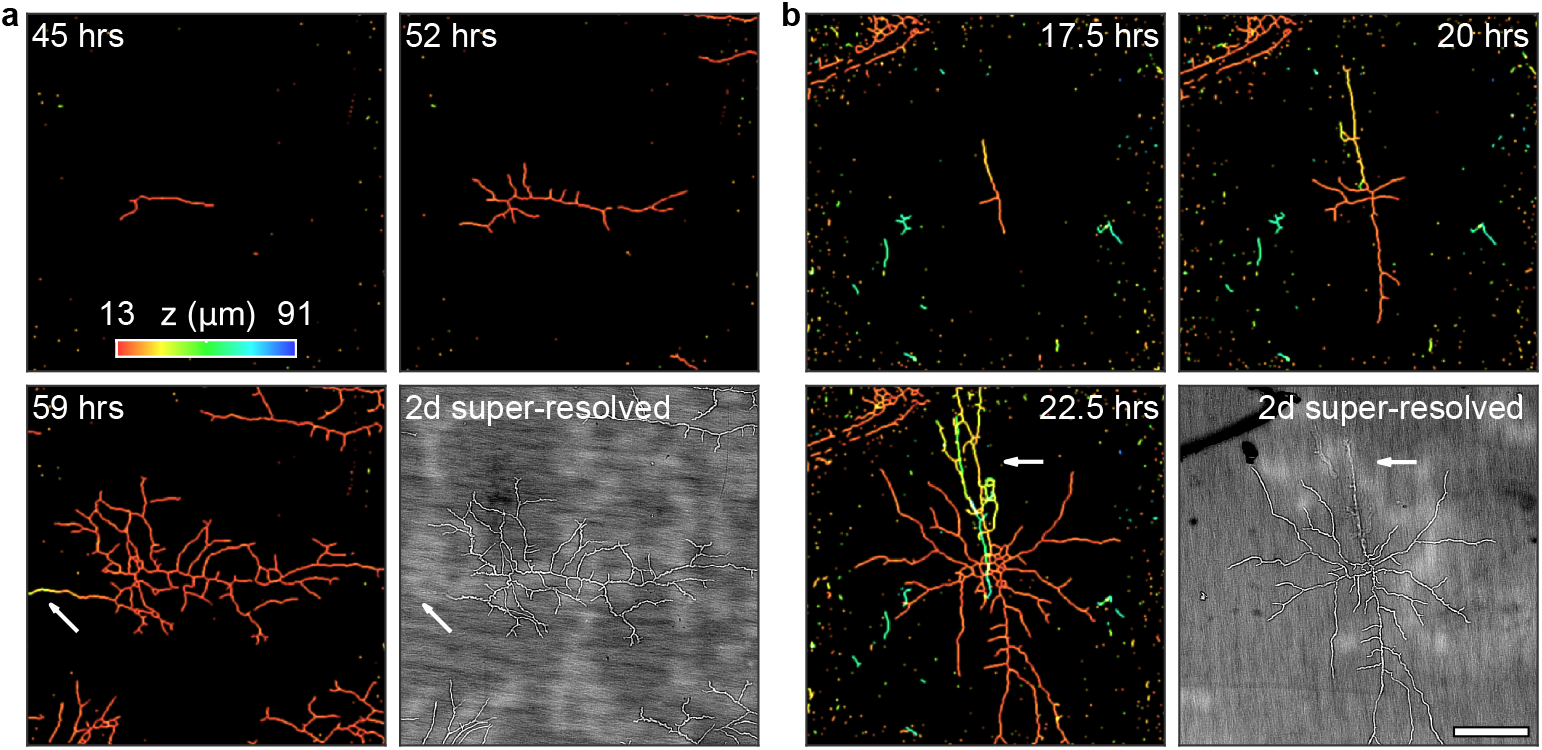
Time-lapse 3D reconstructions and 2D super-resolved images of two captures with filters between *A. brasiliensis* and the growth medium. (a) The majority of hyphae are confined between the filter and optical film. (b) One instance of hyphal penetration through the filter. Scale bar: 200 µm; color bar: height in µm.

## 4. Conclusion

In summary, we introduce an upgraded version of the ePetri dish platform, specifically designed for extended microbial culturing and imaging. Besides hardware changes such as the implementation of a randomized LED array, we emphasize two key innovations. First, the introduction of a sealed growth chamber effectively maintains the growth medium’s integrity for a prolonged period of at least 7 days. This feature is crucial for long-term observations of microorganisms, particularly those that require a stable and consistent growth environment. It is especially beneficial in studies focusing on the growth patterns of bacteria and fungi. Second, the development of a new reconstruction algorithm for the ePetri dish that offers 3D visualization capabilities marks a significant improvement over previous 2D algorithms. The capability of detailed examination of structures such as fungal hyphae makes it particularly valuable in biological studies where 3D structural data is essential, such as in the investigation of complex microbial growth patterns that are unresolvable with 2D imaging.

Looking ahead, one direction to explore is to further refine the 3D reconstruction algorithm. One current limitation is the high video random-access memory (VRAM) requirement due to the extensive number of *z* slices and images processed. We believe that this could be resolved through the application of implicit neural representations [29–31], which might not only address the VRAM issue but also enhance reconstruction quality, especially in densely populated regions. Moreover, the upgraded ePetri dish platform opens new research possibilities in microbiology studies. For example, it allows for the exploration of growth direction preferences in heterogeneous mediums or in response to nutrient sources. Overall, we believe that the ePetri dish will be a powerful platform for studying the intricate and dynamic processes of microbial growth and development, offering unparallel insights into microbiological systems.

## Data Availability

The code and data underlying this study are available from Mango Inc.

